# Coral reefs in crisis: The reliability of deep-time food web reconstructions as analogs for the present

**DOI:** 10.1101/113795

**Authors:** Peter D. Roopnarine, Ashley A. Dineen

## Abstract

Ongoing anthropogenic alterations of the biosphere have shifted emphasis in conservation biology from individual species to entire ecosystems. Modern measures of ecosystem change, however, lack the extended temporal scales necessary to forecast future change under increasingly stressful environmental conditions. Accordingly, the assessment and reconstruction of ecosystem dynamics during previous intervals of environmental stress and climate change in deep time has garnered increasing attention. The nature of the fossil record, though, raises questions about the difficulty of reconstructing paleocommunity and paleoecosystem-level dynamics. In this study, we assess the reliability of such reconstructions by simulating the fossilization of a highly threatened and disturbed modern ecosystem, a Caribbean coral reef. Using a high-resolution coral reef food web from Jamaica, we compare system structures of the modern and simulated fossil reefs, including guild richness and evenness, trophic level distribution, predator dietary breadth, food chain lengths, and modularity. Results indicate that despite the loss of species, guilds, and trophospecies interactions, particularly zooplankton and other soft-bodied organisms, the overall guild diversity, structure, and modularity of the reef ecosystem remained intact. These results have important implications for the integrity of fossil food web studies and coral reef conservation, demonstrating that fossil reef communities can be used to understand reef community dynamics during past regimes of environmental change.

## 1 Introduction

Recent years have witnessed a transition in the emphasis of conservation biology from an initial concern for individual species at risk, to habitat preservation and restoration, and most recently to a focus on entire biological communities and ecosystems [1, 2, 3, 4, 5]. The shift reflects two major developments in conservation biology and community ecology, the first being an acknowledgement that species are parts of larger, integrated systems, themselves part of an integrated biosphere, and that those systems provide services vital to human societies [6]. The nascent field of conservation paleobiology has already incorporated the transition, using multi-taxon approaches as the bases for forecasting models of risk to entire biotas, and paleocommunity models as tools to examine ecosystem dynamics under conditions which lie outside of documented ecological experiences, such as mass extinction [7, 8].

The second development is the growing understanding of communities and ecosystems as complex and rich dynamical structures which often have profound impacts on individual species. Communities have typically been viewed as stable and somewhat static systems (i.e., species interactions are in balanced equilibria) and that stability promotes complexity, but these notions have been replaced by the idea that communities are capable of transitions among multiple, alternative regimes and dynamic equilibria encompassing a range of community parameters [9, 10, 11]. Most recently, it has been suggested that community persistence itself might act as an agent of long-term selection, with functional structures and interactions appearing repeatedly within ecosystems over geological time [12]. Therefore, attempts to understand and conserve modern communities, or the most recent contemporary states of long-lasting communities, must account for the dynamic nature of those communities on multiple timescales and under different environmental circumstances.

The management of modern marine ecosystems undergoing current and future anthropogenically-driven change must be informed by how similar systems have responded to environmental variations in the past. This can be fulfilled at the present time with theoretical and experimental studies, in addition to the examination of the historical and geological records. The latter have the added advantages that the range of environmental conditions to which ecosystems have been subjected in the past is far broader than those within the realm of current scientific and societal experience, and that the possibility of observing alternative dynamic ecosystem regimes is increased when longer intervals of time are considered. For example, a study by Pandolfi and Jackson [13] showed that despite sea level and climate variability, coral reef communities in Barbados were stable in composition for at least 95,000 years during the Pleistocene. Examining and comparing today’s reefs with those of the Pleistocene showed that recent human impacts have resulted in a coral reef structure different from anything seen in the last 220,000 years. Without the establishment of baselines for what non-anthropogenically altered reefs look like, we would not know how altered and degraded today’s marine ecosystems are in comparison [14].

As the future of oceanic ecosystems is still very much uncertain, deep-time studies provide our best proxy for what we can anticipate in the Earth’s near future [15]. For example, major questions in marine conservation and global change biology center on how marine ecosystems will respond to environmental stresses and/or large disturbances, and what makes some communities more vulnerable to perturbation than others [16]. These concerns are driven by the degraded state of ecosystems in today’s oceans, resulting in the decline of species at an alarming rate and unprecedented magnitude [4]. Coral reef ecosystems are of particular interest, because one quarter of all marine species may be found in these threatened marine communities [17, 18, 19, 20].

### 1.1 Preserving the past

One of the foundations of modern paleontology is understanding the impacts of preservation on interpretations of the fossil record, and the ability to use that information to reconstruct the past [21, 22]. Current paleontological studies rarely ignore factors of bias, such as selective preservation among taxa, biased preservation of parts of taxa, and outcrop dimensions. Those concerns must be extended to conservation-based paleobiological approaches. A major complication, however, is that in addition to biases of preservation and discovery, the information conveyed by integrated systems may also be biased by the manner in which integration is preserved. For example, the extent to which prey richness and abundances are preserved surely influences the interpretation of the dynamics of molluscan drilling predation, and differences of generation times bias relative abundances [23, 24]. It is therefore important that we analyze the ways in which fossilization potentially biases our interpretation of community and ecosystem function and ecology. In this chapter we explore the ways in which the retention and loss of data affect our reconstructions of attributes important to measuring system dynamics. Using a high-resolution food web of a modern Jamaican coral reef ecosystem [25], we simulate its fossilization and document subsequent changes of taxon richness and interspecific interactions. We then measure how those changes alter important quantitative measures of food web structure and function, specifically predator dietary breadth, the web’s food chain lengths and trophic level distribution, guild richness and evenness, and the modularity of the system. Finally, we assess the reliability with which we could reconstruct those features of a paleocommunity by assuming that our starting point is the fossilized reef system, then constructing a guild-level representation of the fossil community and contrasting the implied trophic structure to that of a similarly resolved modern reef.

### 1.2 Endangered coral reefs

Threats to modern coral reef ecosystems bear several similarities to conditions from intervals in the Earth’s past, e.g. increasing CO_2_, ocean acidification, and oceanic temperatures. Coral reefs may thus be one of the best proxies we have for predicting future ecological changes and biodiversity loss in the oceans. The ocean is a large sink for anthropogenic carbon dioxide today (~30% of total), and a predicted increase of CO_2_ concentrations in the coming century is expected to adversely affect marine organisms in a multitude of ways, particularly by decreasing biocalcification [26, 27]. Evidence already exists for such a scenario, which is apparent in decreasing calcification rates of individual species and communities, especially in coral reef environments [28]. In addition, global sea surface temperatures (SST) have risen in the past century (0.4-0.8°C), with warming predicted to accelerate in the near future [29]. Increasing temperatures have had a large effect on the marine realm, affecting ocean circulation, benthic and planktonic diversity and abundance, productivity, and overall invertebrate physiology [30, 31]. Coral reefs are extremely sensitive to changes in temperature and pH, frequently expelling their zooxanthellae (photosynthetic algal symbionts) when physiologically stressed, resulting in coral bleaching [18]. During the writing of this chapter (March-April, 2016), Australia’s Great Barrier Reef system is experiencing unprecedented bleaching, with approximately 95% of reefs in the ecosystem being affected [32]. However, corals are not the only organisms under threat in reef ecosystems. Other invertebrates and vertebrates are also in decline due to overfishing, degradation of their coral habitat, and pollution [33, 34, 35, 36, 37, 38].

Caribbean reefs in particular have suffered significant losses, with reports of considerable reduction (~80%) of coral reef cover since the 1970’s in addition to frequent human-induced degradation [39, 40]. Furthermore, in the early 1980’s, a massive disease-induced die-off of the urchin, *Diadema antillarum,* resulted in a macroalgal bloom that persists to this day [41, 42, 43]. The less biodiverse algal-dominated state of Jamaica’s reefs in particular is exacerbated by historical overexploitation of herbivorous fish [37]. It appears, however, that *Diadema* has been functionally replaced to some extent by parrotfishes, highlighting the potential importance of functional redundancy in coral reef and other ecosystems [42, 44, 37, 45]. Regardless, local increases of parrotfish in various areas of the Caribbean have not reversed the algal-dominated state of the reefs, although studies have indicated that when parrotfish are able to escape overfishing inside marine reserves, they were able to increase grazing intensity and reduce macroalgal cover [46]. Recent reports do show a slight recovery of *Diadema* and a subsequent increase in coral recruitment and survivorship, though not nearly as rapidly as expected, with current populations at only 11.62% of their premortem density [47,48]. Jamaican reefs have come to represent an unhealthy reef system, frequently disturbed by anthropogenic (e.g., overfishing and pollution) and non-anthropogenic (i.e., disease, hurricanes) stresses [49].

Today climate change and other human-induced disturbances are progressing at an unparalleled and alarming rate, increasing the need for studies that use novel analogues and proxies from Earth’s past. Data from the fossil record give us a longterm historical perspective from which we can test the influence of extreme environmental conditions on ecological dynamics and community structure. Fossil food web studies are particularly needed due to recent studies indicating that functional extinction of apex predators, large herbivores, or ecosystem engineers in coastal ecosystems may occur several decades to centuries after the onset of ecosystem degradation, resulting in potential collapse of trophic webs [34, 50]. Thus, deeptime studies have much to contribute to evaluating potential losses of biodiversity, stability, and sustainability in marine ecosystems due to current and future climate change. For example, Aronson [51] determined that Antarctic benthic food web structure was established 41 million years ago when the climate was much cooler, resulting in predators (i.e., sharks, crabs, etc.) being pushed from Antarctic waters. An increase in current temperatures as a result of climate change may result in the the invasion of such durophagous predators, profoundly affecting benthic food web structure. As such, paleoecological perspectives are vital to modern conservation strategies in order to establish how we can buffer anthropogenic and climate-related stress in fragile modern ecosystems currently and in the future.

## 2 Fossilizing a coral reef

The Jamaican coral reef food web describes species interactions from coral reef and adjacent seagrass habitats within Jamaica’s marine geopolitical territory, including the offshore Pedro Bank (Fig. 1). The food web model is an amalgamation of data drawn from multiple specific localities. Differences between localities, however, are expected to be ephemeral and changing; this is supported by the significant compositional overlap both among localities within Jamaica, as well as with similar food webs constructed for the neighbouring Cayman Islands and Cuba [25]. This northern Caribbean region represents a common regional species pool, and the Jamaican dataset is thus a sample of that pool, integrated over the spatial variation present among the Jamaican localities. Taxon composition was determined from published compilations and reports, including Fishbase [52], and the REEF [53] survey database compiled up to 2011. Details of sources and methods used to determine species interactions are given in [25], and the complete data are archived in the DRYAD database [54]. Together, the resources represent approximately 50 years of data, a temporal resolution which is likely much finer than any available in the fossil record. Comparison to sub-fossil and archaeological data from Jamaica, however, suggest that compositionally the data would be congruent with fossil data time-averaged on at least a millennial scale [55]. The complete dataset documents the interactions of 749 species in the northern Caribbean, ranging from single-celled protists to multicellular macrophytes and metazoans, of which 728 have records of occurrence in Jamaica. Multiple species were collapsed into trophospecies when they shared exactly the same prey and predators (i.e., had exactly the same interactions in the food web). This process resulted in 265 trophospecies, of which 249 are present in Jamaica, with a total of 4,105 inter-trophospecific predator-prey interactions.

**Fig. 1.**
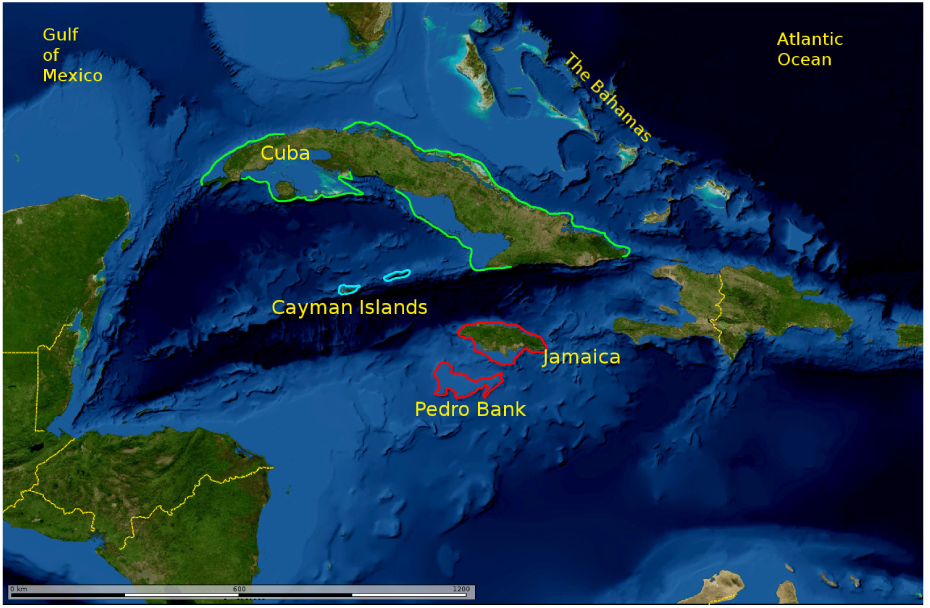
Map of the Greater Antilles, showing the regions covered by the coral reef data (adapted from [25]). Data treated in this study are from Jamaican geopolitical areas, outlined in red.

Fossilization of the community was simulated with a simple binary filter at the genus level. The occurrence of the genus to which each taxon in the food web is assigned was checked for occurrence in the fossil record using the Paleobiology Database (downloaded September, 2014) and Sepkoski’s Compendium of Fossil Marine Genera [56], and was considered fossilized if the genus has a documented fossil record. The presumption is based on the premise that characteristics which promote fossilization, such as morphology, life habits and habitat, can be extended to all members of the genus. If the genus does not have a documented fossil record, then the food web taxon was eliminated from the dataset. This simulated fossilization resulted in a reduced dataset comprising 433 species aggregated into 172 trophospecies, and 1,737 inter-trophospecific interactions. We tested whether the likelihood of species fossilization is uniform among the trophospecies, suspecting that biases would exist because of the usual vagaries of fossilization, including biases against soft-bodied taxa, small body size, easily disarticulated skeletons, and depositional environment. The expected number of species fossilized in a trophospecies is estimated simply as the species richness of the trophospecies times the overall fraction of preservation for all the trophospecies,

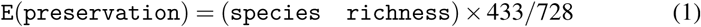

Thus the probability that the number of species lost (i.e., not fossilized) from a single trophospecies is consistent with uniform probabilities of non-preservation among all trophospecies, is given by the hypergeometric probability

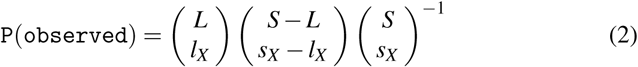

where *S* is food web species richness, *L* is the number of species lost during simulated fosilization, *s_X_* is the richness of trophospecies *X*, and *l_X_* is species loss from *X* during fossilization. A hypothesis of uniform, unbiased levels of preservation could not be rejected for 130 trophospecies. The remaining trophospecies had either improbably low or high levels of preservation (Fig. 2), and all can be explained by body composition and body size. For example, trophospecies with unexpectedly poor preservation include nanno-zooplankton (p = 0.0006), epibenthic sponges (p = 1.02 × 10^−8^), micro-zooplankton trophospecies such as cyclopoid copepods (p = 0.0003), gorgonians (p = 8.91 × 10^−6^), and sphenopid and zoanthid anthozoans (p = 8.91 × 10^−6^). In contrast, trophospecies with unexpectedly good preservation include mixotrophic scleractinian corals (p = 3.95 × 10^−15^) and soft-sediment dwelling, infaunal suspension feeding bivalves (p = 0.0001).

**Fig. 2.**
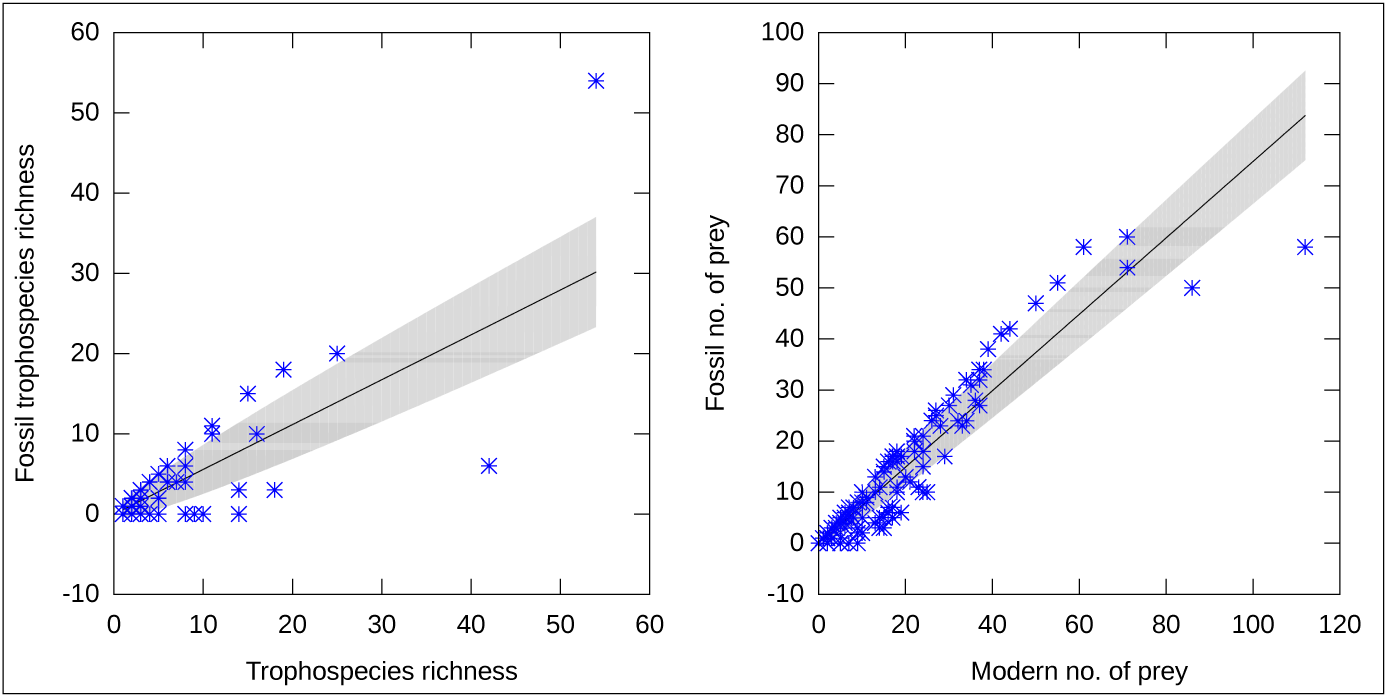
Expected and observed losses of taxa (left) and interactions (right). Values after simulated fossilization are plotted against the modern values. The expected values, based on an assumption of a uniform probability of preservation among all trophospecies, is plotted as a line. Grey regions represent 1.96 standard deviations around the expected values (see Appendix 1).

A loss of biotic interactions is expected to accompany the loss of taxa with lower probabilities of preservation. We examined the distribution of lost interactions among the trophospecies, expecting that the loss would be a function of the number of interactions of a trophospecies; that is, more connected trophospecies would lose a proportionally greater number of interactions. Biased preservation of species within some trophospecies, however, would also render other trophospecies more poorly connected, or disconnected, than expected based on their numbers of interactions alone. We therefore examined incoming, that is, predatory interactions for each trophospecies using a hypergeometric probability similar to that explained above for taxon preservation. Given that 1,737 trophospecies interactions out of 4,105 were preserved, the expected number of interactions (prey) retained by a trophospecies is estimated as

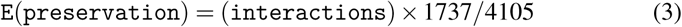

The probability of observing the number actually lost is then

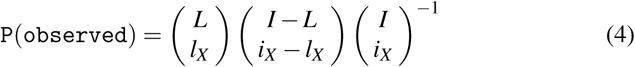

where *I* is the total number of interactions in the food web, *i_X_* is the number of incoming interactions, or prey, of trophospecies *X*, and *L* is the total number of interactions lost. Trophospecies that lost more incoming interactions (i.e., predatory interactions) than expected include scleractinian corals (p = 0.003 for both mixotrophic and fully heterotrophic trophospecies), the butterflyfish *Chaetodon capistratus* (p = 0.004), and a multi-taxon trophospecies including the fish blue chromis *(Chromis cyanea),* brown chromis (C. *multilineata)* and royal gramma (*Gramma loreto*) (p = 0.003). Zooplankton constitute a major or sometimes total portion of the diet of all these taxa, and the significantly low preservation probabilities of zooplankton trophospecies result in their predators being either poorly connected or completely disconnected from the fossil food web. In contrast, a number of fish trophospecies retain more interactions than expected, primarily because benthic invertebrates with hard body-parts, and hence high probabilities of preservation, dominate their diets. This set of consumers includes the seabream (*Diplodus caudimaculatus*) (p = 0.00007), the pufferfish (*Sphoeroides spengleri*) (p = 3.1 x 10^−7^) and the squirrelfish (*Sargocentron vexillarium*) (p = 0.00004).

The reconstruction of taxon level trophic properties is obviously affected by the differential probabilities of preservation of taxa, based primarily on body composition and body size, and consequently the preservation of trophic interactions. The low preservation of a major group such as the copepod zooplankton not only generates a negative bias against the inferred dietary breadths of their predators, but also creates a positive bias toward taxa whose prey have exceptionally high probabilities of preservation. These results are not surprising given what we know about the vagaries and biases of fossil preservation, but the relevant question here is what are the biases created when viewing the community as an integrated system, and not merely as a collection of taxa and interactions. We therefore examined several system-level properties, including the distribution of dietary breadths (“in-degree” distribution), food chain lengths, trophic levels, modularity, and guild structure and diversity.

### 2.1 Dietary breadth

The distribution of dietary breadths is the distribution of the number of prey species per consumer species, or in the case of aggregated food webs such as this, the number of trophospecies preyed upon by each consumer trophospecies. This distribution is commonly referred to in the food web literature as an in-degree distribution, where degree refers to the number of interactions per species, trophospecies, or trophic guild. Surveys of these distributions [57, 58, 59] show that the overwhelming majority are “decay” distributions, where the density of the distribution is biased toward low dietary breadths, meaning that there are more taxa with specialized diets in the web than there are species with generalist diets. The precise nature of a web’s distribution is unknowable unless all interactions have been recorded, but estimates from a variety of webs and models suggest that the distributions may be exponential, power law, or a mixture of decay distributions [60]. One interesting feature is that the tails of these distributions are long (hyperbolic) and occupied by generalist species with broad diets. Topological analyses of food web networks suggest that the presence of such highly connected species provides community robustness against the cascading effects of random extinctions, because those taxa are less likely to lose all their connections and thus provide some insurance to the community [61]. That conclusion must be tempered, however, by a corresponding weakness of many of those links, because a general theoretical conclusion is that strong interactions destabilize species interactions and hence communities in general [62, 63, 64]. Furthermore, empirical measures of interaction strengths between modern species show that the majority of those interactions are relatively weak [65]. Reconstructions of Lagerstatten food webs, with presumably high probabilities of preservation, have also yielded hyperbolic distributions [66]. Models of paleocommunity food web dynamics have used hyperbolic decay distributions, that is, fat-tailed distributions, but the majority of interactions in the community models are typically weak [67, 68, 12].

Given the above observations of taxon and interaction loss during fossilization, however, can paleocommunities be reliable sources for the reconstruction of indegree distributions? A comparison of the modern and simulated fossil distributions for the coral reef community shows that, at least in this case, a reconstruction based on the paleocommunity would be very accurate (Fig. 3). The modern distribution can be described best with a power function, ln(*p*) *=* 3.48 – 0.78[ln(*r*)] where *r* is the number of prey trophospecies per consumer trophospecies, that is consumer in-degree, and *p* is the number of consumer trophospecies of that degree (F(1, 54) = 130.74, r^2^ = 0.702, p < 0.0001). The fossil distribution is essentially identical, ln(p) = 3.05 — 0.78[ln(r)], (F(1,37) = 95.41, r^2^ = 0.713, p < 0.0001), and statistically indistinguishable (Student’s t = 0.01, p = 0.990). Thus, despite the loss of taxa and interactions, including the structurally and functionally important zooplankton trophospecies, one can predict the number of interactions per fossil trophospecies with a high degree of confidence.

**Fig. 3.**
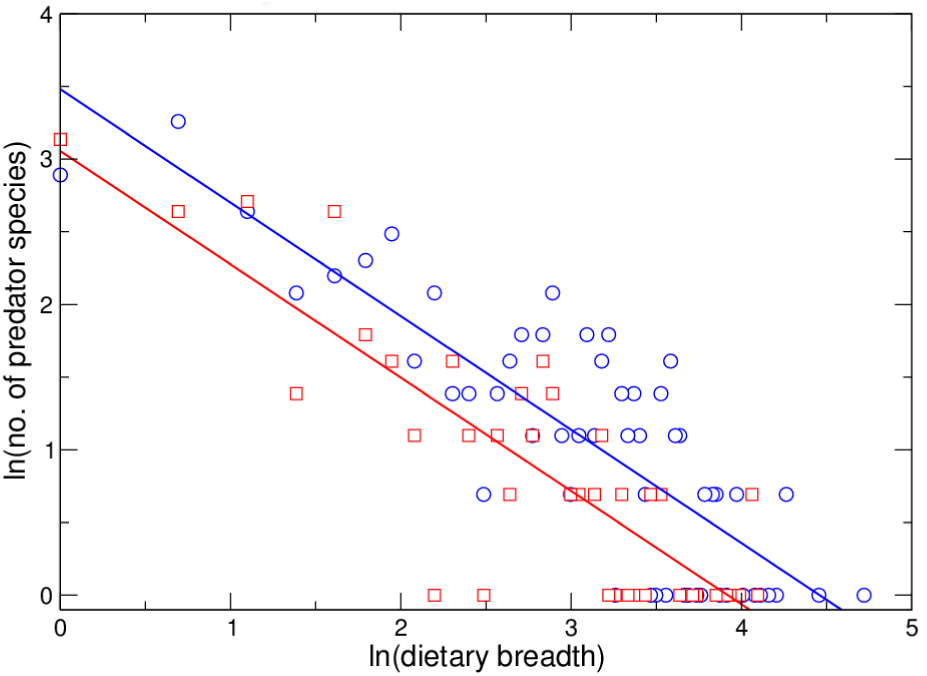
Trophic in-degree, or dietary breadth, distributions. The number of predatory species with a specific number of prey or resource species (y-axis) is plotted against that specific number of prey, (x-axis). Both the modern (open circles) and fossil (open squares) distributions are fit significantly by logarithmic power functions.

### 2.2 Trophic chains and levels

Trophic level is an important characteristic in both species and system conservation efforts [69, 70, 71]. It is generally accepted that high trophic level species, that is, predators with superior capabilities and few predators of their own, are often at greater risk from anthropogenic actions. The reasons for this vary between marine and terrestrial habitats. In the former, high trophic level species are often targeted for harvesting because of high biomass, while terrestrially they are viewed as threats to domestic livestock or competitors for resources, and are often sensitive to habitat alteration or destruction. The impact of overfishing on oceanic predators has been devastating, and populations throughout the Caribbean are in decline [34, 37]. Scleractinian reefs, however, originated and evolved during a series of changing predatory clades, including Mesozoic marine reptiles, and the radiations of modern chondrichthyans, teleosts, and teleost clades specialized to reef environments [72]. The potential richness and complexity of coral reef systems might therefore vary with the relative proportions of different life history strategies present, and different suites of trophic levels would represent alternative, yet persistent ecosystem regimes.

Trophic level is broadly understood to describe, in some manner, the position of a species within a food web, if species are arranged hierarchically from basal, primary producers, up to apex predators. Species positions in the food web determine the flow of energy to their populations, and the feasibility of their persistence depends on a productive basal component, sufficient to support all those populations further up the food chains [73]. Energy is lost thermodynamically along each step of a food chain, however, because of respiration and the inefficiency of energy assimilation from consumed material, and this loss is thought to constrain the lengths of food chains [74]. Food chain length may also be constrained by the likelihood of decreasing dynamical stability as the length of a chain grows, [75], and by numerous factors specific to a particular food web, its composition and environmental context [76].

All these limitations would restrict the diversity and complexity of predation within a community, but there is variability driven by organismal variation. Yodzis, thinking in terms of webs rather than chains, suggested that increased productivity would make both increased chain length and predator diversity feasible [77]. This in turn means that any predator toward the top of a food chain would have to be a super-generalist, spreading its effort over a greater diversity of prey, in order to garner sufficient energy. This effort could be focused on other predators near the top, or by omnivory, where the predator would feed at multiple levels of the chain thus circumventing some of the thermodynamic loss. Finally Pimm (1982), noting the decreasing trend of efficiency of energy conversion moving from ectothermic invertebrates, to ectothermic vertebrates, to endothermic vertebrates, suggested that invertebrates should support longer chains [78]. The generally larger population sizes of smaller species, and greater rates of invertebrate population growth, would also increase the likelihood of dynamical stability [76].

Variability of complexity is driven in modern Caribbean reefs largely by anthropogenic factors [34, 79], but over longer timescales this variability could be a function of clade diversity dynamics and macroevolutionary trends. Identification of any such trends, and the establishment of reliable historical baselines for modern reefs, depends on whether trophic chains and levels can be reconstructed from fossil and sub-fossil data. As such, here we introduce a method for calculating trophic position from food web network data, network trophic level, and use it to compare the distribution of trophic levels in the modern and fossilized reefs. The network trophic level (ntl) of a species or trophospecies is the average shortest distance of its prey species to a primary producer. Primary producers are assigned an ntl of 1.0, and the ntl of a consumer species i is calculated as

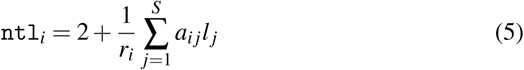

where *r_i_* is the number of prey species of *i*, *a_ij_ =* 1 if *i* preys on species *j,* and zero otherwise, and *l_j_* is the shortest path length of *j* to a primary producer. Ntl differs from the prey-averaged trophic level of Williams and Martinez [73] by a factor of one for consumers. Trophic level is measured in various ways, ranging from simplistic integer values corresponding to discrete categories such as “primary”, “secondary”, “tertiary” consumer and so forth, to inferences of the number of steps to a consumer as measured by stable isotopic composition of consumer tissues. A common measure used for fish is fractional trophic level (ftl). Ftl is based on the proportion of specific prey species in a consumer’s diet, and is a weighted average of the prey ftl values. Romanuk et al. [80] presented a global database of empirically measured and inferred ftl values for fish. Using that database, Roopnarine [81] showed that ftl and ntl are correlated significantly (Fig. 4), concluding that ntl is a reliable measure of trophic level, and that a significant proportion of trophic level variance is based only on position in the food web.

**Fig. 4.**
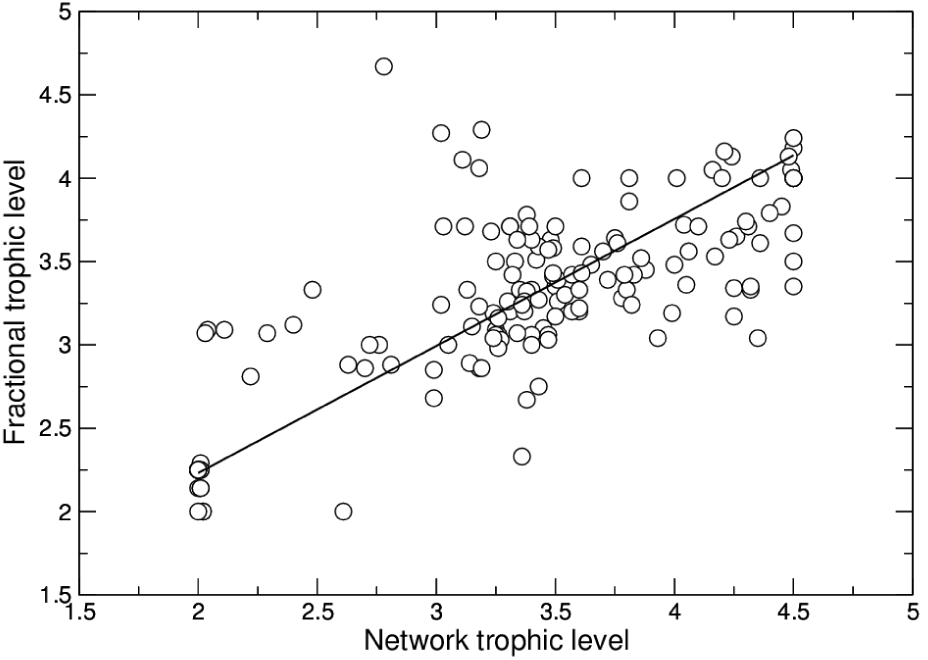
The relationship between network trophic level and fractional trophic level, fit with a reduced major axis function.

Consumer ntl ranges from 2-5.5 in the Jamaican reef food web, with a mean value of 2.89 which increases to 3.29 if primary consumers (ntl = 2) are excluded. Almost all trophospecies with ntl values between 4 and 5 are vertebrates, primarily predatory fish with a broad range of body sizes. Carnivorous ophiuroids is the single invertebrate trophospecies in this range, the included species being both benthic deposit feeders and polychaete carnivores (e.g. *Ophiocoma echinata*). All trophic levels of five and above, however, are occupied by invertebrate trophospecies, specifically corallivorous polychaetes and gastropods, e.g. *Hermodice carunculata* and *Coral-liophila caribbaea* (ntl = 5.0), and gastropod predators of polychaetes, e.g. *Conus regius* (ntl = 5.5). These very high trophic levels are the result of very long food chains which extend the phytoplankton-zooplankton food chain, but which are also very simple, meaning that the prey breadths of trophospecies along the chain tend to be low. The high ntl invertebrate taxa are also not apex predators, but instead are subject to predation by trophospecies that feed at multiple trophic levels and along multiple food chains. For example, *C. regius* is preyed upon by a trophospecies of carnivorous crustaceans, including *Penaeus duorarum,* with a ntl of 3.18. The assessment of trophic level is therefore complicated by the branching topology of a food web. Ntl, as perhaps with other trophic level measures such as stable isotopic composition [76], is thus not a linear ordination of taxa among multiple food chains. They are, however, reliable measures of the average distances of consumers from the productive base of a web. The ntl measures reported here are consistent with other observations that invertebrates tend to occupy longer food chains than vertebrates [82, 74, 83]. Moreover, although high ntl vertebrate predators frequently prey on invertebrates along long food chains, those vertebrates also feed in much shorter food chains and thus again have lower ntl values. For example, the top predator Caribbean reef shark *Carcharhinusperezi* is of ntl 3.86. The fact that the most powerful predators in a food web network will not be the furthest removed from primary producers is a warning against the common practice of simplifying food web structures into discrete, or nearly discrete trophic levels.

The ntl distribution of the modern reef is significantly higher than the simulated fossil reef (Fig. 5; Kruskal-Wallis, *χ*^2^ *=* 30.70, p = 0.0001). Maximum ntl in the modern reef is 5.5, but only 4 in the fossil reef. The reduction could be the result of a loss of high ntl trophospecies, but there is no significant difference between the ntl distributions of those trophospecies that are preserved and those that have no fossil representation (Student’s t = 1.612, p = 0.108). The difference of ntl distribution is in fact attributable to the reduction of ntl of preserved trophospecies. Many of those trophospecies are specialized predators with poorly preserved prey, such as carnivorous gastropod predators of gorgonians or polychaetes, or zooplanktivorous fish. Those consumers’ ntl values have collapsed to two, implying incorrectly that they are primary consumers. Other trophospecies ntl values are reduced because they have no preserved prey and their ntl values are 1.0. Thus, the poor preservation of key trophospecies such as zooplankton and soft-bodied taxa has a significant impact on estimates of fossil trophic level, and the impact will always lead to an underestimation of trophic level and food chain lengths. There is essentially no way in which data can be recovered to address this issue, and the evolution and historical baselines of trophospecies trophic level in coral reef ecosystems, and indeed all paleoecosystems, are obscured by biases of preservation. It is conceivable, however, that information is retained, and can be inferred, at higher levels of system organization that might be less subject to bias, and we turn to those in the following sections.

**Fig. 5.**
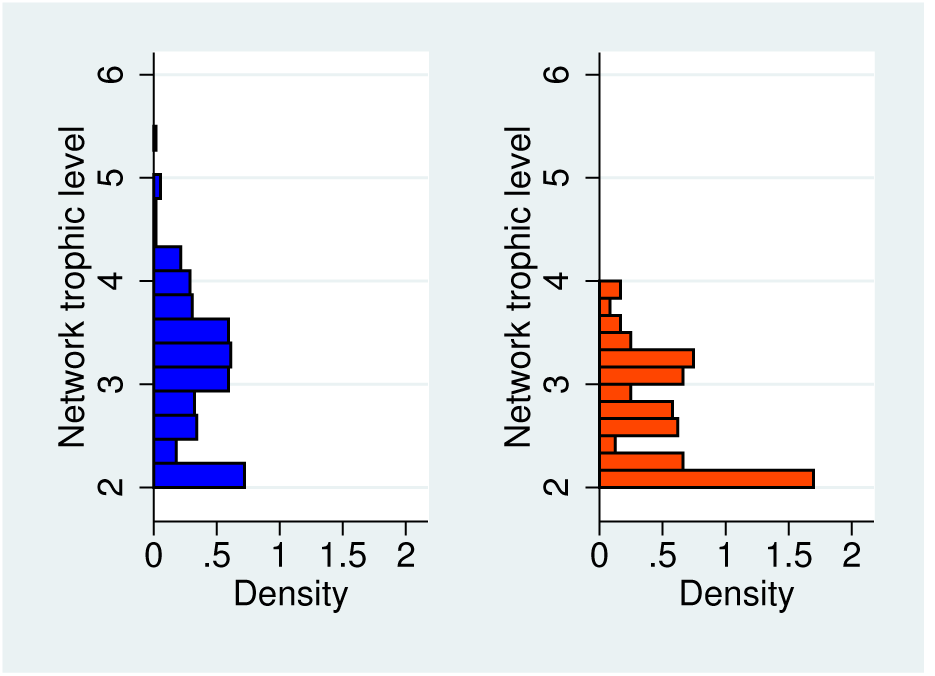
Network trophic level (ntl) distributions for the modern (left) and simulated fossil (right) food webs. The fossil distribution is truncated, with a maximum ntl value of 4 in contrast to the modern web’s 5.5. Primary producers (ntl=1) are excluded from the plots.

### 2.3 Modularity

A module is a subset of nodes within a network which have more interactions with each other, and fewer with other nodes, than would be expected if interactions occurred at random [84]. Obvious examples occur in social networks where persons within a family or circle of friends may represent a module within the larger society.

A module within a food web would therefore comprise species sharing more interactions with each other than they do with other species in the community, a condition also referred to in the food web literature as compartmentalization. In offering this definition, we distinguish this usage of “module” and “modularity” from an alternative use which refers to pairs or trios of interacting species without regard to their other interspecific interactions (e.g., [85]. We also discount trivial compartmentalizations that result in discrete, non-overlapping sets of interactions, for example as might occur across strong habitat boundaries. Such compartments are in effect independent food webs. We focus instead on compartments embedded in, and sharing interactions with the rest of the network.

Interest in food web modularity stems from May’s theoretical work [62] on the relationship between the local stability of a community (i.e., its ability to return to a static equilibrium after a minor perturbation), and the community’s richness, connectance (the density of interspecific interactions), and average interaction strength. May noted that for random networks or food webs, the probability of stability decreases with an increase of any of those community parameters, calling into question the long-held hypothesis of a positive relationship between community complexity and stability. May also pointed toward possible “solutions” to this seeming paradox, including a hypothesis that many food webs might consist of compartments, or modules, which would increase the probability of stability. We suggest here that in addition to stable dynamics, modular structure of a food web would indicate ways in which the energy supplied to a community is partitioned, and the extent to which the community could then be viewed as energetically integrated or compartmentalized.

Results of subsequent searches for food web modularity have been mostly equivocal [86], possibly because of the relatively low resolution of many current food web datasets. Another probable cause is the difficulty of identifying modules in highly resolved, complex food webs. In a community of *S* species, the number of modules could range from 1 (all species within a single module) to S (each species represents a separate module). All modular arrangements of size between these two extremes would represent all possible combinations of partitioning schemes of S, and that number grows very rapidly with increasing *S*! There are no objective and exact analytical approaches, short of an exhaustive search of all those possible combinations of species clusters. Heuristic approaches do exist, however, and here we applied the modularity algorithm due to Newman [87], which is an optimization of the property Q, where Q = (fraction of interactions within modules) - (expected fraction of such interactions). We used a fast approximation of Newman’s algorithm developed by Blondel et al. [88], as implemented in the network visualization software Gephi [89], to generate repeated modularity measures on the same network. We further tested the null hypothesis that the community network is indistinguishable from a random network of equal species richness and number of interactions, by comparing the food web to such equivalent random networks using the netcarto program [90, 91, 92].

Both the netcarto and Blondel et al. implementations support the sub-division of the food web into four modules (Fig. 6), a relative modularity value of 0.282, and with the netcarto randomizations failing to support the null hypothesis of no unique modularity (z-test; mean relative modularity of 1,000 random networks = 0.139, sd = 0.002, p < 0.0001). The coral reef community thus comprises four modules or sub-communities, and that substructure is highly unlikely to arise by any random assembly of species and interactions. The collapse of 249 trophospecies into four modules is striking, but do the modules make any sense ecologically? Our expectation that the food web would be partitioned among different conduits of energy flow is partially correct, but factors of habitat and modes of life also play a role. Figure 7 illustrates summaries of the modules, presented as food chains and featuring examples of the trophospecies within each module.

**Fig. 6.**
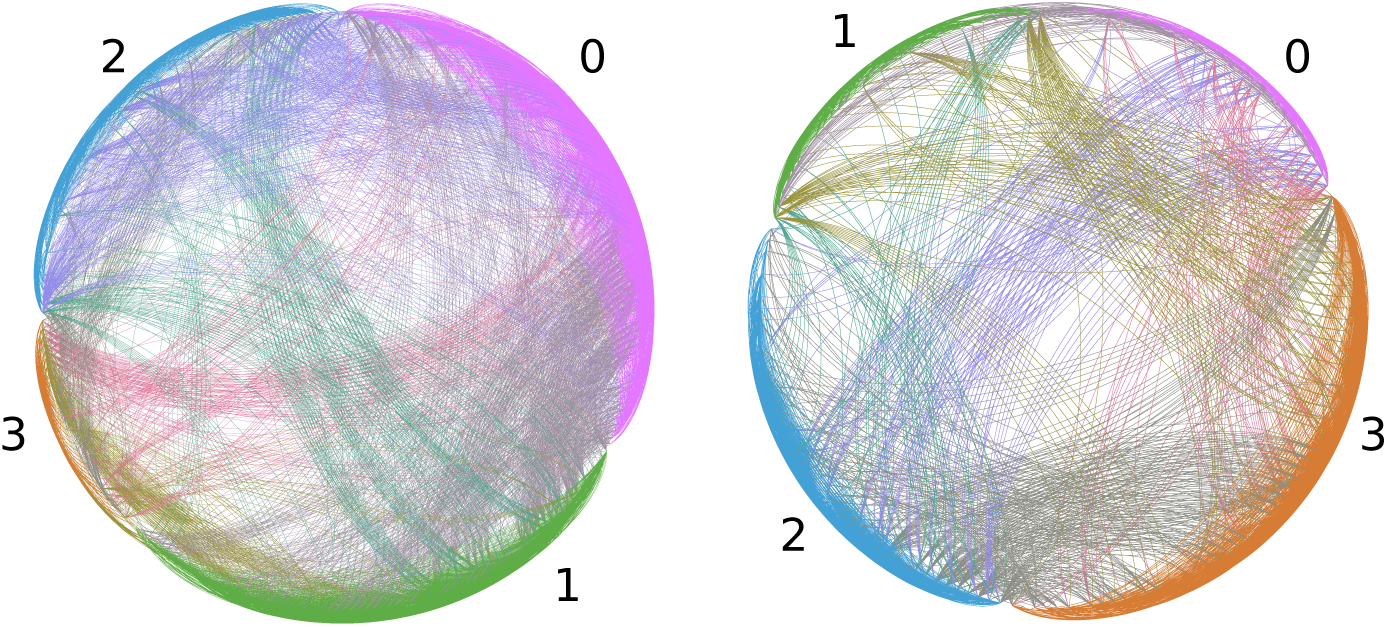
Modularity of the modern (left) and simulated fossil (right) food webs. Numbers indicating each module are explained in the text, and correspond to the same module between the webs. Visually, modules are identifiable by the density of within-module interactions which are shown on the periphery, in contrast to between-module interactions which cross the interior of the circularly arranged webs.

**Fig. 7.**
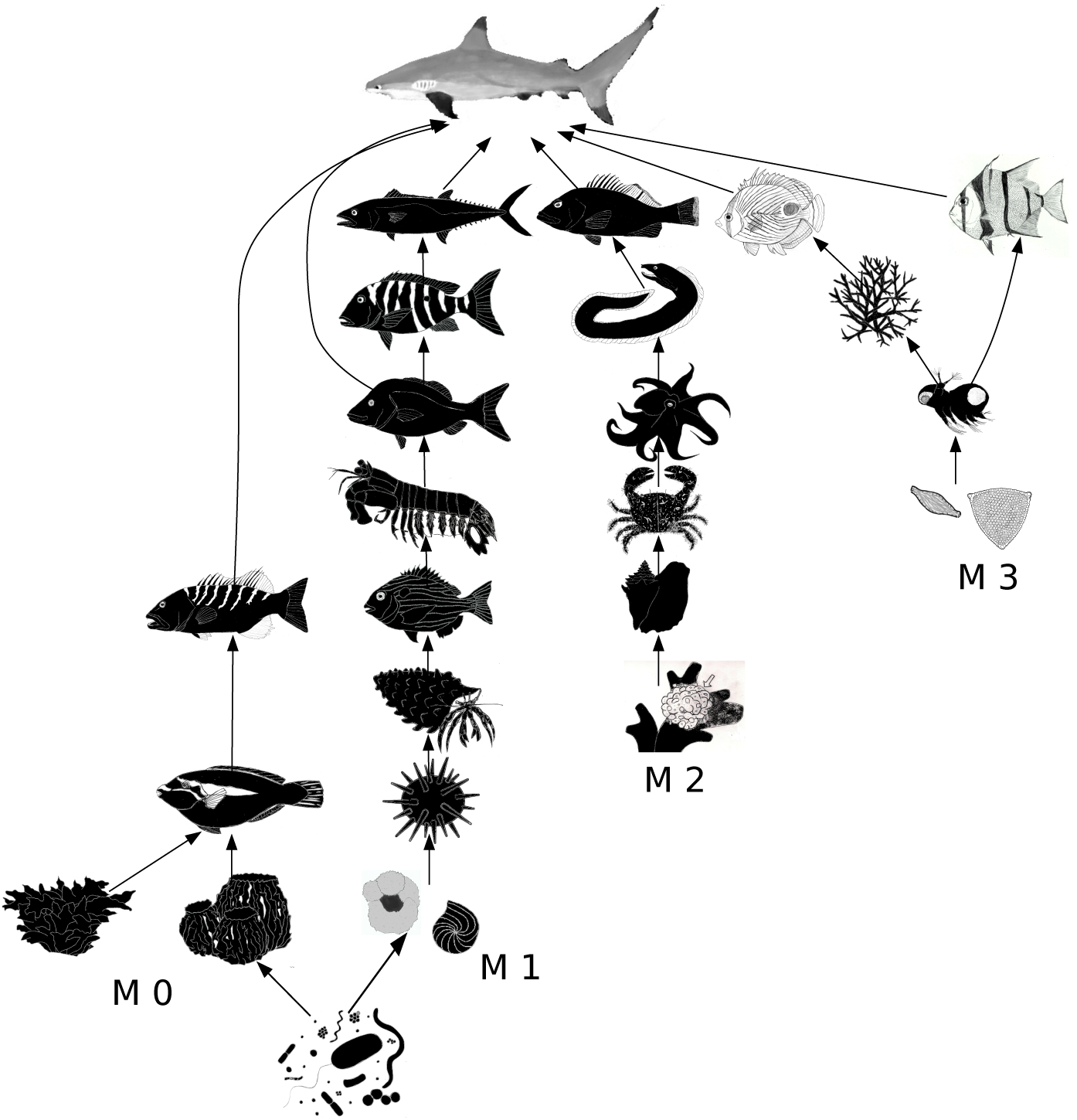
Representative food chains within each of the modern community’s modules. From left to right, trophospecies, or representative species, selected from each module are: M0 - bacteria, the sheet macroalga *Ulva,* epibenthic basket sponge *Xetospongia,* parrot fish *Scarus iserti,* tiger grouper *Mycteroperca tigris,* and Caribbean reef shark *Carcharhinus perezi*; M1 - foraminifera *Globigerina* and *Archaias,* the sea urchin *Lytechinus,* hermit crab *Pagurus brevidactylus,* seabream *Archosargus rhomboidalis,* mantis shrimp *Gonodactylus,* margate *Haemulon album,* mutton snapper *Lutjanus analis,* and king mackerel *Scomberomorus cavalla;* M2 - coralline algae *Hydrolithon,* queen conch *Strombus gigas,* hairy crab *Pilumnus marshi,* reef octopus *Octopus briareus,* purple-mouth moray *Gymnothorax vicinus,* and red hind *Epinephelus guttatus;* M3 - epiphytic diatoms, copepods *(Podon* sp. shown), staghorn coral *Acropora cervicornis,* reef butterflyfish *Chaetodon capistratus*, and Atlantic spadefish *Chaetodipterus faber*.

Module 0 (modules are numbered here 0, 1, 2 and 3), composed of 34 trophospecies, is a basal energetic module in the sense that it includes all the macrophytic trophospecies, including macroalgae and seagrasses. It also includes the sponges and herbivorous fish as well as low trophic level omnivores that consume macroalgae and benthic sponges, such as the parrot fish *Scarus iserti.* The highest trophic level trophospecies in the module includes piscivores that specialize on benthic grazing fish, e.g. the tiger grouper *Mycteroperca tigris,* and generalist predatory piscivores, including the apex predator Caribbean reef shark *Carcharhinus perezi.*

Module 1, comprising 66 trophospecies, is dominated by benthic food chains, including benthic foraminifera and metazoan deposit feeders, and grazers such as lytechinid echinoids. Food chains are extended through the module primarily by omnivorous and carnivorous benthic macrocrustaceans, e.g. pagurid crabs, and both benthic invertebrate predators and teleost predators of those taxa, including gonodactylid stomatopods. The food chains are again capped within the module by high trophic level piscivores such as the king mackerel *Scomberomorus cavalla.*

Module 2 does not have a notable primary producer base, containing only encrusting coralline algae, but the module size is nevertheless substantial, comprising 74 trophospecies. Much of this module’s diversity is dominated by benthic macroinvertebrates, including herbivorous, omnivorous and carnivorous grazers, such as the queen conch *Strombus gigas,* the hairy crab *Pilumnus marshi,* and the reef octopus *Octopus briareus* respectively. Higher trophic level predators in the module are primarily fish specialized to predation in the coral habitat, such as moray eels and their predators.

Module 3, a plankton-based module, is the largest module with 75 trophospecies, and includes the major phytoplankton and zooplankton trophospecies, as well as major benthic and pelagic zooplanktivores, such as corals and the Atlantic spadefish *Chaetodipterus faber.* There are fewer high trophic level predators in the module, and its trophospecies richness is a function of the great diversity of trophic strategies employed by zooplankton and zooplanktivores.

Although the community is modular, and the modules are highly interpretable in ecological terms, the modules are united by high trophic level predators, such as the Caribbean reef shark. This predator, though assigned to a single module, preys on species in all modules. An examination of other large shark species in the northern Caribbean region, as documented in similar food webs from the Cayman Islands and Cuba [25], reveals similar broad, generalist predation. However, most of those species have been exterminated on local scales by overfishing and are rare in regional reef systems today. Prior to their extirpation, modularity of the Jamaican reef would have been weaker because of greater high trophic level, cross-module predation. Therefore, in spite of the theoretical expectations of a positive role for modularity, measurements based on modern, anthropogenically-altered communities might result in overestimates of natural modularity. This possibility could be addressed with historical and paleontological records, but only if modularity is preserved in spite of the losses of taxa and interactions. We therefore repeated the modularity analyses using the simulated fossilized community.

Modularity analysis of the fossilized food web yielded four modules, and a modularity measure of 0.287. Comparison to 1,000 random networks failed to support a null hypothesis of insignificant modularity (z-test; mean relative modularity of random networks = 0.169, sd = 0.002, p < 0.0001). The number of modules equals that for the modern food web, and the modularities are very similar. The important question though is how comparable are the modules in the two webs? To answer this, we compared the membership of preserved trophospecies in both modern and fossil modules, examining the distribution of fossil modules within each modern module (Fig. 8). There is a single dominant fossil module occurring in each modern module, suggesting significantly that those fossil modules are equivalent to the modern modules (Chi-square test, *χ*^2^ *=* 273.606, p < 0.0001). Accepting this hypothesis results in only 13 fossil trophospecies not corresponding to their presumed modern modules, a correct classification rate of 92.4%. Remarkably, despite the loss of trophospecies and interactions, the modular structure of the modern reef is expected to be preserved in fossil reef communities.

**Fig. 8.**
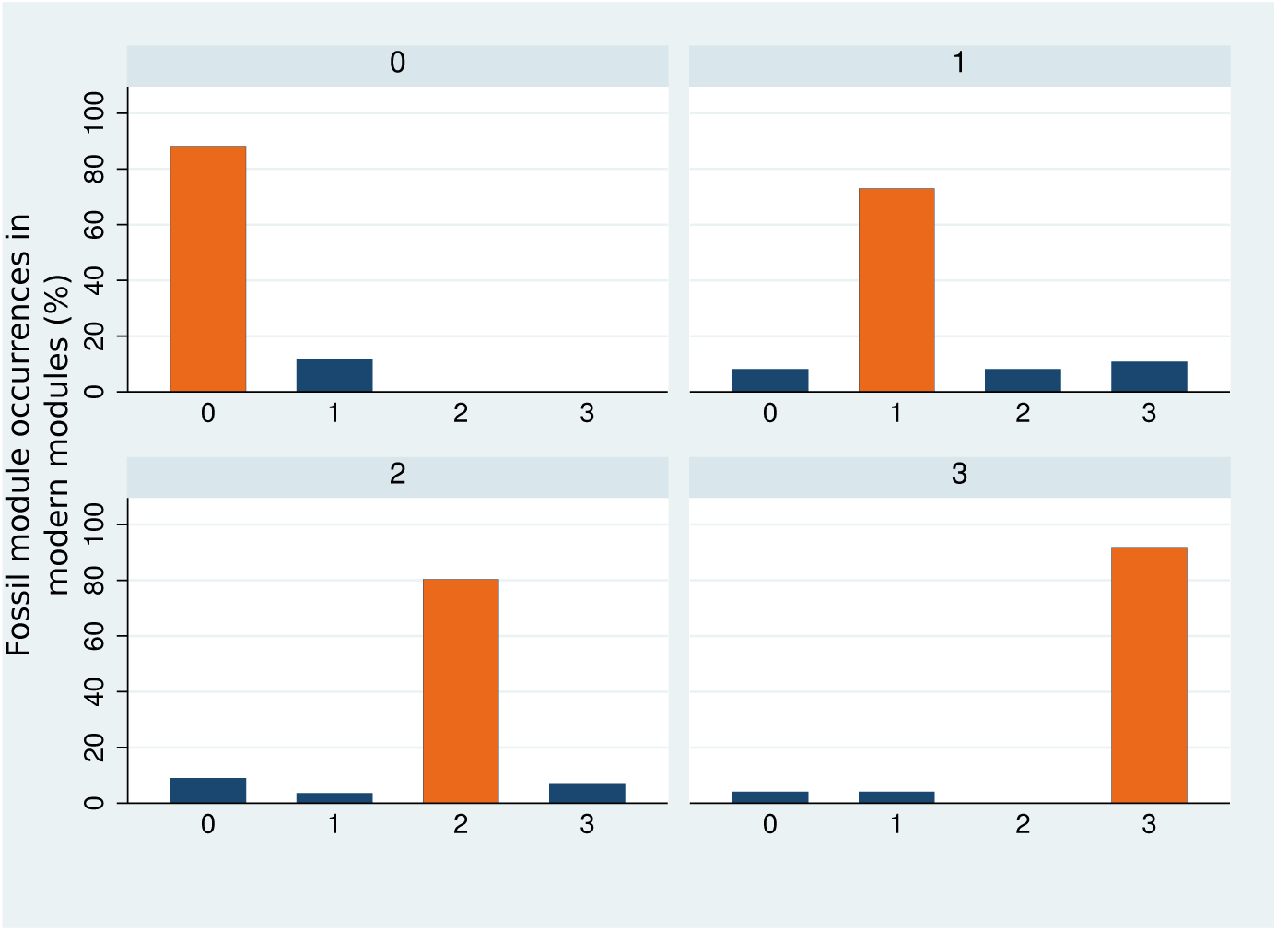
The distribution of modules identified in the simulated fossil food web, within each module of the modern food web. The statistically dominant occurrence of a single fossil module in each modern module supports the equivalency of the webs’ modular structures. Each modern module is plotted separately, as indicated at the top of each plot. Modules in the fossil food web are shown on x-axes, numbered 0-3, to indicate significant statistical equivalence to their modern counterparts.

## 3 Guild structure and diversity

Food web reconstructions often aggregate species into groups on the basis of presumed ecological similarities, such as “benthic macroinvertebrates”, “salt marsh plants”, etc. Aggregation is perhaps justified when more precise data are not available, or more precise data are unsuitable for the analytical models being employed. The extent to which aggregation obscures or distorts food web structure and dynamics is an unsettled argument, but one solution would be to simply increase the precision with which food web networks are constructed. A version of this solution employs the trophospecies, or “trophic species” approach, in which species aggregates comprise taxa with exactly the same trophic interactions [93]. Trophospecies simplify a food web without sacrificing precision, and is the approach taken in the construction of the modern Caribbean coral reef community. The trophospecies approach has also been applied to the fossil record in at least three instances, including reconstructions of Cambrian marine food webs of the Burgess Shale and Chengjiang faunas [66], and an Eocene terrestrial food web of the Messel Shale [94]. All these reconstructions take advantage of fossil Lagerstätte, where fossil preservation is exquisite, probabilities of preservation are high, and some traces of trophic interactions are available. Furthermore, although taxon preservation is almost certainly incomplete, the Messel Shale food web includes 700 taxa, and with the current coral reef food web represents one of the largest food web reconstructions available. Thus, in spite of incomplete preservation, fossil food webs are amongst the best food web reconstructions currently available. There are three drawbacks to the Lagerstätten approach however, and those are that one is limited to the time, place and community type of the Lagerstätten. Additionally, Cambrian marine food webs are unlikely to offer much insight into the dynamics of coral reef ecosystems under conditions of a rapidly changing ocean, as the majority of Cambrian taxa are not extant.

Another solution to the problem of aggregation is to estimate precise interspecific interactions even when data on interactions are limited. Simply put, this approach models the precision required for a species-level food web by estimating the number of interactions per species and the species which interact, while constraining those estimates on the basis of known or inferred ecologies. This approach has been developed most extensively in analyses of Paleozoic and Mesozoic terrestrial food webs around the end Permian and end Cretaceous mass extinctions [67, 12]. Those analyses used very well preserved biotas from the Karoo Basin of South Africa, and western North America respectively, but their preservation falls below Lagerstätten quality. Roopnarine and co-workers addressed this problem by assigning taxa to trophic guilds, which are aggregations based on common body size, habitat, and potentially overlapping prey and predators. Coral reef examples would include the placement of all carcharhinid sharks into a guild, or all epibenthic sponges into a guild. Trophic interactions between guilds lack the precision of interspecific interactions, but they are in actuality sets of interspecific interactions, and each interspecific interaction belongs to a single interguild interaction. Given a community of species, there is a finite number of food webs that can be constructed, but many of them would not be consistent with the ecologies and interactions of the community. Guild level food webs, also termed metanetworks, limit this number to a subset of species level food webs that are consistent with ecological reality. The dynamic terrestrial models of Roopnarine et al. [67, 12] sample species level food webs from the metanetwork subset.

An unresolved question is how well guilds and metanetworks actually capture the functional structure of a community. We address this here by first identifying the guild structure of the reef, and then taking the perspective that one is required to reconstruct the guild structure given only those taxa that would be preserved in the simulated fossil reef. The reliability of the fossil metanetwork is then evaluated by comparing its diversity and evenness to the modern metanetwork, and examining whether the reef’s distribution of trophic levels can be reconstructed from the fossil metanetwork.

### 3.1 Identifying guilds in a food web

The purpose of a trophic guild is to aggregate species with overlapping prey and predators. Trophospecies are therefore a type of guild, one in which members share exactly the same prey and predators. Membership can be extended, while maintaining exact overlap, to species with different numbers of prey and predators, but where the interactions of some of the species are subsets of others, i.e., the interactions of one species are nested within those of another. For example, prey of the porcupine fish *Chilomycterus antennatus* is a subset of the trophically more general trigger-fish, *Batistes vetula.* A requirement of exact overlap, however, becomes problematic under two conditions. First, increasingly precise data are liable to uncover small differences between species, thereby separating species which were formerly assigned to the same trophospecies. Second, if most interactions are indeed relatively weak, then how much weight should be assigned to interactions that are not shared between species, versus those that are shared? In both cases species can be ecologically similar, and yet their aggregation in a food web be ambiguous. The questions could potentially be answered for modern communities by ever-increasing empirical work, although these are very difficult data to obtain. Furthermore, the questions simply cannot be answered for paleocommunities. Here we propose a heuristic solution where we use the overlap of interactions among species to recognize guilds, but limit the expected overlap according to the limits of fossil preservation.

The procedure begins with a pairwise measure of interaction overlap between all species in the food web. Overlap is measured separately for both types of interactions, predator-prey and prey-predator, as

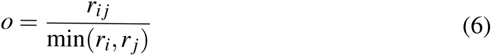

where *r_i_* is the number of prey (predators) of species *i*, and *r_ij_* is the number of prey (predators) that *i* and *j* have in common. The value of *o* is thus zero if the species have no prey in common, and 1 if they have all prey in common or one of the prey sets is nested completely within the other. Predator overlap is measured similarly, counting number or predators for *r_i_* and *r_ij_*, such that o equals zero when the species have no predators in common. Each set of measures yields a *S* × *S* matrix of overlap measures, where S is the species richness of the community. The two matrices are then combined for each pair of species as the simple product of prey and predator overlap, producing a single matrix of overlap indices, **O**. The resulting elements of **O** then range from 0, where either prey or predator interactions, or both, fail to overlap between the two species, to 1, where overlap or nestedness is complete. Within the **O** matrix will be groups of species which overlap amongst themselves more strongly than they do with other species, in effect forming modules. Guilds can therefore be identified by examining the modularity of the **O** matrix, and would be equivalent to **O**’s modules.

Given the concerns expressed above regarding the precision of overlap, we examined the matrix’s modularity, and hence guild composition, at multiple levels of overlap by applying thresholds, where values below a threshold would be excluded from guild recognition. We proceeded from a threshold value of 0.1, where species sharing an overlap greater than 0.1 could potentially be assigned to the same guild, to 1, where species within a guild would have perfectly nested sets of interactions. Guild inclusivity thus decreases as the threshold increases, and the number of guilds should therefore increase. These expectations are indeed met, as the number of guilds in the coral reef community, and the strength of overlap as measured by Newman’s modularity index, increase as the threshold increases, with modularity being at a maximum when the threshold is 0.9 (Fig. 9). The number of guilds increases from 10 at a threshold of 0.1, to 149 at a threshold of 1. The 728 species in the Jamaican coral reef food web, represented by 249 trophospecies, may therefore be nested within 149 guilds. This number is comparable to work bycBambach [95], in which 118 modes of life were found to exist for recent marine metazoans. Given the greater resolution available in our data, 118 most likely represents a lower limit of the number of guilds.

**Fig. 9.**
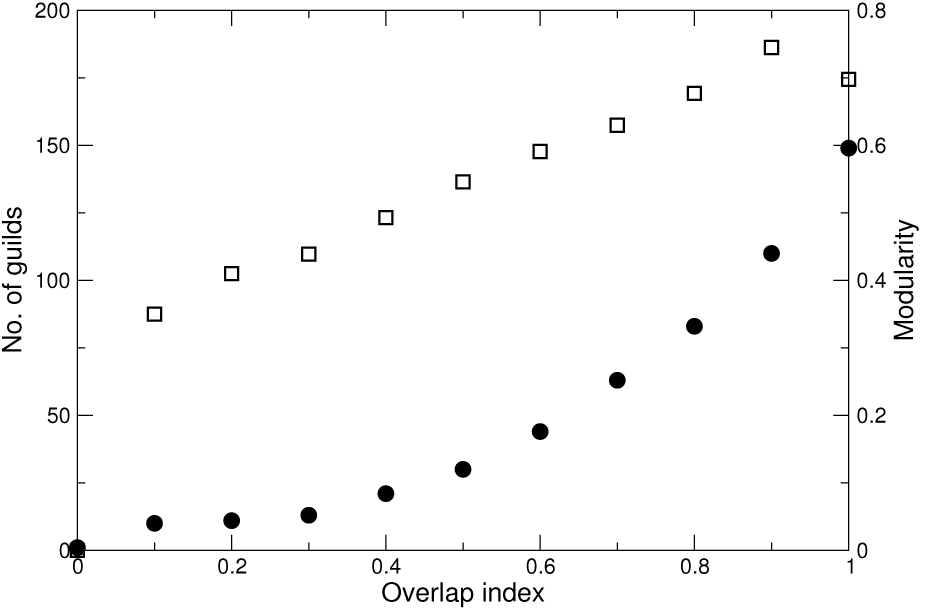
The number of guilds, or modules, recognized as the overlap or nestedness of interactions between species. The extent of overlap, or threshold, is indicated on the x axis, and the corresponding number of guilds (filled circles), and strength of the modularity (open squares), are indicated on the left and y axes respectively.

## 4 Reconstructing the community

Imagine that our starting point is not a modern coral reef, but a fossil coral reef instead. Can an understanding of how community information is transformed during fossilization be combined with fossil taxon richness to reconstruct a paleocommunity food web? The easiest reconstruction is at the guild level, where species are aggregated into sets of overlapping interactions. Not all the 149 guilds of the modern reef would be preserved, however, and many interactions are also lost. The losses cannot be accounted for in many cases, for example the absence of a guild of corallivorous polychaetes, because not only do those species lack a fossil record, but it is doubtful that their existence could otherwise be inferred from preserved taxa. There are two rich and highly connected groups, however, whose existence can be inferred even though they are absent from the fossil record: epibenthic sponges and zooplankton. The sponges do not have a taxonomic record, but their presence is recorded as large numbers of disarticulated spicules. The sponges are a key link between pelagic microorganisms on which they prey and guilds of spongivorous macroinvertebrates and vertebrates. Major groups within the zooplankton guilds are also absent from the fossil record, e.g copepods, or have no records which include coral reef-dwelling members, e.g. scyphozoans. There are large numbers of preserved zooplanktivorous species and guilds, however, such as corals and many other cnidarians, and zooplanktivorous fish. We could therefore insert guilds representing the missing groups into the set of preserved guilds with defensible confidence, and connect them to appropriate prey and predator guilds; but two problems must be addressed.

First, whereas the epibenthic sponge trophospecies is assigned to a single guild out of 149 in the modern food web, micro-zooplankton are distributed among six different guilds, and macro-zooplankton are assigned to two guilds, one of which also contains micro-zooplankton. Given the absence of zooplankton from our fossil data, we would have no idea how many zooplankton guilds should be inserted into the reconstructed community. The best and most conservative answer would be to use a threshold of overlap at which a single zooplankton guild first emerges, thereby aggregating all zooplankton into a single guild, yet distinguishing them from other guilds. We therefore examined guild structure at each threshold as described above (Fig. 9), and the zooplankton first emerge as a single guild at the 0.6 level as thresholds increase from 0.1 to 1. There are 42 guilds at this threshold (see Appendix 2, Table 1), and we consider it to be the best resolution that could be inferred from our knowledge of fossil taxa, in the absence of modern data. Thirty three of the 42 guilds include fossilizable taxa.

Second, how many species should be assigned to the sponge and zooplankton guilds? Deriving sponge richness from spicule diversity is notoriously difficult and unreliable [96], and the zooplankton essentially have no fossil record in the coral reef system. We addressed this problem by referring to a similar situation in terrestrial paleocommunities with insect faunas. Mitchell et al. [68], in their reconstructions of Late Cretaceous North American communities, estimated insect richness based on a positive relationship between the insect richness of well-preserved faunas spanning the Phanerozoic, and the richness of vertebrate predation on those faunas. The relationship is consistent over this time interval. The logic of the relationship is that predator and prey richnesses may often be related in communities over evolutionary time, since a greater diversity of prey could support a greater diversity of predators, both energetically and by reducing competition among the predators. Similarly, the presence of predators generally supports a greater diversity of prey species. We therefore examined this relationship for a zooplankton food chain at the 0.6 threshold. Guilds on this food chain include “phytoplankton”, “nanno-zooplankton”, “foraminifera” and other heterotrophic protists, “zooplankton”, “gorgonians” and “pelagic planktivores”. The set of observations consists of the richness of a prey guild, and the sum of predators supported by that guild, weighted by number of prey guilds linked to each predatory guild. That is, predator richness

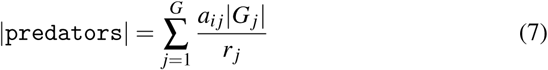

where | G_*j*_ | is richness of a guild of predators linked to prey guild *i*, and *r_j_* is the number of guilds preyed upon by predatory guild *j.* The relationship was fit using reduced major axis (RMA) regression both because neither variable can be considered independent, and neither is measured without error (Fig. 10). Furthermore, a RMA function is symmetrical, meaning that the function is invariant to the choice of which variable is treated as the predictor. The RMA function,

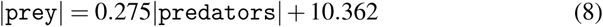

is significant (r^2^ = 0.97, F = 270.468, p = 0.00005), and predicts a zooplankton guild richness of 45 species. This is in excellent agreement with the actual richness of zooplankton species (44).

**Fig. 10.**
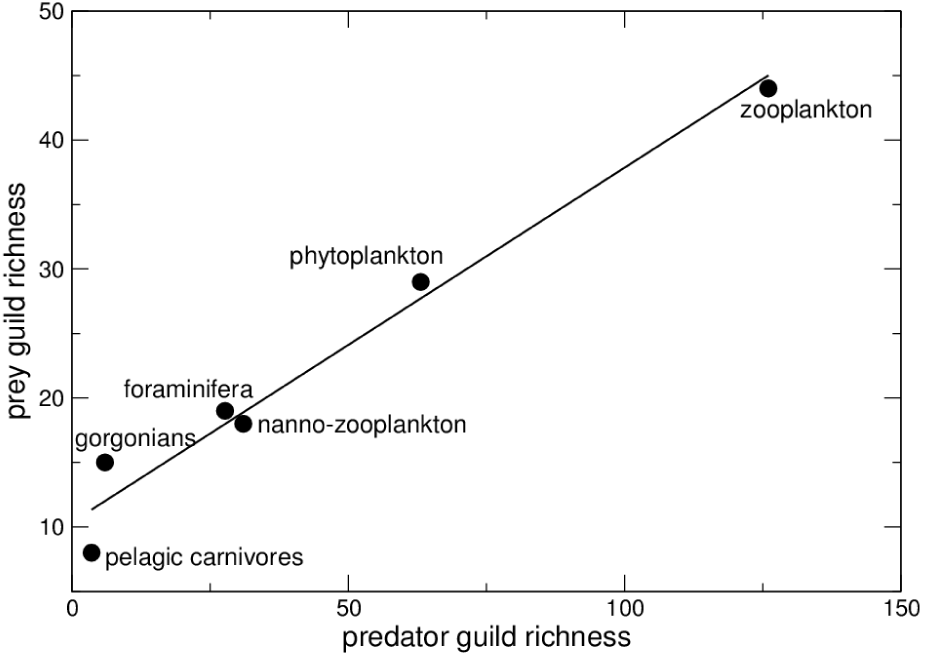
The relationship between predator guild richness and prey guild richness for guilds along the major “zooplankton” food chain. The line indicates a fitted reduced major axis function. Prey guilds are indicated at each corresponding data point.

A similar function could not be constructed successfully for the sponges, because the sponge guild-level food chain at 42 guild resolution includes two highly aggregated guilds, the “hard benthic macroinvertebrates” and “large vertebrate macrophyte and invertebrate grazers” which confound any more specific numerical relationships between prey and predator richnesses. The overall fraction of species preserved in the simulated paleocommunity is 0.59 (433 out of 728 species). Sponge richness was therefore estimated using the fraction of overall preservation (0.59), yielding (0.59 x 53) = 31 species, which is greater than the 16 species (all endolithic) expected to be fossilized, but closer to the original richness of 53 species.

### 4.1 Diversity and evenness

In order to explore the similarities and/or differences between the modern and fossil coral reef food webs, we adopted methods commonly employed in paleoecology, calculating the Simpson Index of Diversity (1-D) and the Shannon Index (H’) to examine the richness and distribution of species across guilds [97, 98]. The Simpson Index of Diversity (1-D) is a common metric for quantifying taxonomic diversity and abundance, and is used here to estimate the taxonomic richness of guilds in the modern and fossil coral reef communities. Hence, the higher the value of the Simpsons Index of Diversity, the greater the taxonomic diversity of the guilds. The Shannon Index (H’) is used to calculate the evenness 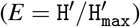 of the distribution of species across guilds, with a maximum value of 1.0 indicating a perfectly even distribution and a minimum value of 0 indicating a highly uneven distribution.

In applying these metrics to the modern Jamaican coral reef community, consisting of 728 species within 42 guilds, we found that the community had very high taxonomic diversity, with a Simpson Index of 0.92. Meanwhile, the fossilized coral reef community, consisting of 33 guilds and 433 species, had a very similar taxonomic diversity, with a Simpson Index of 0.88. The modern coral reef community was also fairly even in its distribution of species across guilds, with an evenness (E) value of 0.75; the fossil reef community was comparable with E= 0.72. In addition, as previously discussed, one can safely assume via fossil evidence (e.g., sponge spicules) that a zooplankton and a sponge guild are likely to be present. By replenishing those two guilds (34 guilds present, 493 species) and recalculating the diversity metrics, we found that the community now had a Simpson Index (1-D) of 0.90 and an evenness (E) value of 0.74. As such, it is apparent that by restoring these unfossilized taxa to the fossil community, the taxonomic diversity values become almost identical to that of the modern community.

However, while it is reassuring that these two diversity metrics are comparable between the modern and fossilized communities, we have the foresight to be aware that ten guilds that appear in the modern coral reef community are not preserved in the fossilized community. Thus, it would appear that caution must be taken in relying on these metrics to quantify guild structure; other metrics and analyses, such as in-degree distribution, modularity, and trophic levels are needed in order to fully characterise food web structure. Nonetheless, these metrics show that both the modern and fossil coral reef communities had high taxonomic richness, though potentially lacking some redundancy within guilds. These results agree with previous hypotheses indicating that coral reef ecosystems, despite containing a high diversity of species and guilds, are extremely vulnerable to environmental and anthropogenic perturbations given the potential for the loss of key ecosystem processes and decreased resilience due to an uneven distribution of species across guilds [99, 100, 101, 102, 45]. Low redundancy within guilds, as implied here, would further exacerbate this vulnerability.

### 4.2 Simulated food webs

A primary goal of paleo-food web reconstruction has been the modelling of paleocommunity and ecosystem dynamics [67, 68, 12, 103, 104, 59]. Those models require the simulation of food webs that are consistent with paleocommunity structure, as described by resolvable guild-level networks. The reconstruction of the coral reef at the guild level presented here shows that guild structure is preserved during the simulated fossilization, within limits set by the absence of data on at least two significant groups, the zooplankton and epibenthic sponges. Furthermore, the data can be improved with reliable estimates of species for those missing guilds, with analyses of diversity and evenness supporting the improvement. A concluding step in this exercise is then the derivation of a species-level food web from the preserved guild structure. We did this by assigning in-degrees (number of prey) to each consumer species with random draws from the in-degree equation derived above (Section 2.1), constrained to the total richness of the guilds upon which the species could potentially prey. The actual prey species are then assigned randomly to the consumer (see [59] for further details). The resulting simulated food web consists of 398 connected species, with 965 interactions. The latter number is far below the 4,105 inter-trophospecific interactions of the modern community, but recall that the simulated food web is based on an aggregation into 44 guilds. A second food web was simulated, but in this case using a mixed exponential-power law indegree distribution used commonly in previous paleo-food web dynamics studies [67, 68, 12, 103, 104]. This distribution takes the form *P*(*r*) = *e*^−*r*/*ε*^, where *r* is the in-degree of the consumer, *ε =* e^(*γ* − 1) in(*Μ*)/(*γ*)^, *M* is the total number of prey potentially available to a consumer, and γ, the power law exponent, is 2.5. The value of the exponent is the mid-point of a range previously explored for this distribution when applied to Permian-Triassic paleocommunities from the Karoo Basin of South Africa [67]. Distributions within the range were found to construct species-level food webs with linkage densities and connectances comparable to those reported for modern food webs [105, 106]. Applying the distribution to the system of fossil coral reef guilds here, yields a web with a greater density of interactions, with 8,924 interspecific interactions.

It is conceivable that, despite the disparities of interactions among the webs, they could nevertheless yield similar dynamical properties. Such similarity would be possible, however, only if the patterns of interaction are similar among the webs [12]. This was checked for the species-level fossil and modern webs by comparing their network trophic level (ntl) distributions. Recall that those distributions differed significantly between the modern and simulated fossil webs (Fig. 5). The question addressed here is whether the reconstructions, based upon the reconstruction of guild structure, and restoration of unfossilized zooplankton and epibenthic sponges, mitigates any of that loss of structure. The ntl distribution of the simulated web based on the observed in-degree power law distribution differs significantly from the modern web (ANOVA; df = 2,478; F = 30.25, p < 0.0001), and a majority of taxa are below ntl 3.0 (Fig. 11). The ntl distribution of the web based on the mixed exponential-power law distribution, however, is statistically indistinguishable from the modern web (Scheffe’s multiple comparison test, p = 1.0), and reconstructs both the mean (3.29 in both cases) and maximum values (5.5 and 6 respectively). The mixed exponential-power law distribution predicts denser interactions than does the power law, in effect being “fatter-tailed”. This feature compensates for the loss of interactions due to the loss of taxa, and obviously reconstructs the hierarchical structure of the modern food web with significant fidelity. We predict that dynamic analyses of fossil coral reef communities based on this type of reconstruction would be comparable to analyses of modern communities.

**Fig. 11.**
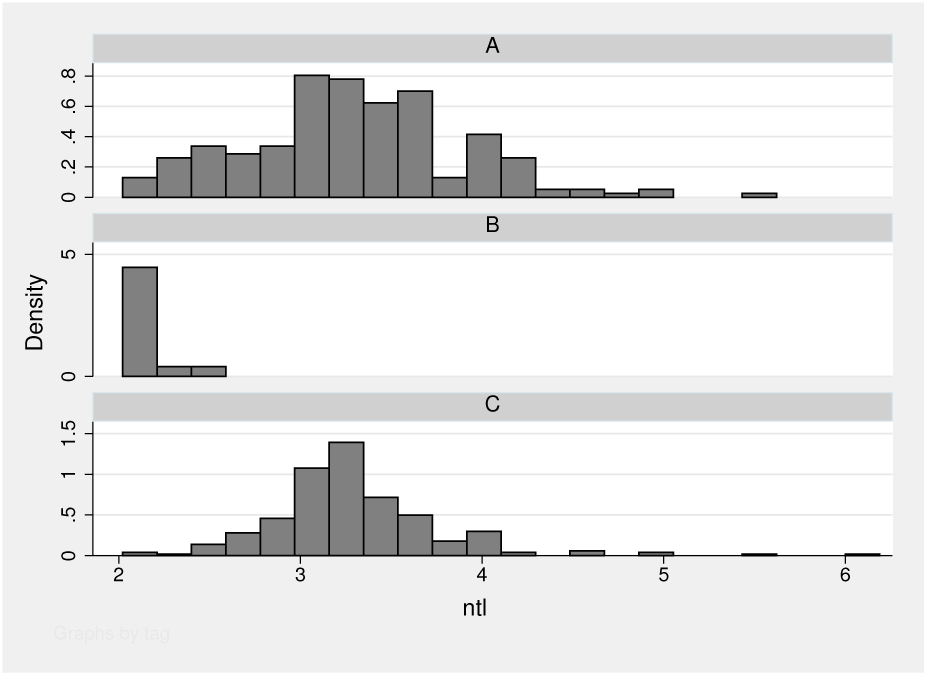
Observed and reconstructed network trophic level (ntl) distributions for all secondary and greater consumers. A - observed distribution measured from the modern food web. B - The ntl distribution based on a species-level food web reconstructed from the fossil guild network, and using the power law in-degree (dietary breadth) distribution of the modern food web. C - A similar reconstruction, but made using a mixed exponential-power law in-degree function. The power law-based reconstruction in B fails to recreate the trophic level distribution, yielding a highly truncated and short-chained food web. The mixed distribution produces a food web with trophic level distribution statistically indistinguishable from the modern food web.

## 5 Summary

The ongoing, unprecedented, global, anthropogenically-driven degradation of natural systems has led to a need for conservation strategies that account for whole ecosystem structures. Coral reefs have received much attention in regards to the effects of anthropogenic and non-anthropogenic stressors on ecosystem functioning, as recent surveys have indicated a global decline in reef diversity, coral cover, and overall functioning [19, 107]. Caribbean reefs, in particular, have suffered considerable historical damage and overexploitation, potentially since initial colonization of islands and coastal areas by humans [37]. Studies of modern coral reef functioning and diversity cannot capture the shifted baselines of species composition, population sizes, and community structure, although these are likely required to forecast what we can expect in a future of continuing environmental decline. Deep time studies have the potential to compliment modern studies by providing insight into how prior coral reef systems have responded to similar environmental perturbations, or perturbations of similar magnitude, in Earth’s past [108].

Here, we tested the plausibility for such studies by simulating the fossilization of a modern Jamaican coral reef food web, and determining how realistically a food web could be recreated from the fossil record [25]. The initial modern Jamaican food web consisted of 728 species, which were then collapsed into 249 trophospecies (i.e., species that share exactly the same prey and predators) with a total of 4,105 inter-trophospecific predator-prey interactions. Simulated fossilization, or removal of genera without a documented fossil record, resulted in a community comprising 433 species, 172 trophospecies and 1,737 inter-trophospecific interactions. As expected, the loss of trophospecies and interspecific interactions resulted in a significant bias, both against poorly preserved species, and in favour of those species that had better than average fossil records. Particularly significant was the almost complete loss of zooplankton trophospecies, which resulted in their predators being either poorly connected or completely disconnected from the fossil food web. While the absence of zooplankton in the simulated fossil data places a negative bias against zooplanktivorous species in terms of biotic interactions, the importance of taxa whose prey have exceptionally high probabilities of preservation, e.g. durophagous crustaceans and fish, may be overestimated.

A positive consequence of the latter bias, coupled with the preservation of more than half of the species in the food web, is that several important features are retained by the simulated fossil community, namely the distribution of dietary breadths among consumers, and the modularity or compartmentalization of the community. The distribution of dietary breadths, or in-degree distribution, describes the number of resource species per consumer species. Compilations based on a variety of modern food webs suggest that those distributions are generally of a decay type and hyperbolic. This means that more species are specialists, consuming relatively few species, and fewer species are generalists; yet the hyperbolic nature of the distributions also implies that generalist species occur at frequencies greater than would be expected were dietary breadths distributed normally. The coral reef meets these expectations, being fit significantly with a power function. The distribution for the reduced, simulated fossilized web is statistically indistinguishable from the modern web’s distribution. Nevertheless, the loss of interactions does result in a significant alteration of the trophic level structure of the community. Many species have lower trophic levels in the fossil community, particularly those involved in very long food chains that include taxa with low probabilities of preservation. The result is that the distribution of trophic levels in the simulated fossil community is significantly truncated, and there is no possibility of recreating trophic levels based on taxon composition alone.

Modularity has been proposed to be important to the stability or resistance of food webs to perturbations, yet the search for modules in food webs has yielded equivocal results. Here we demonstrated that the Jamaican coral reef food web is indeed modular. Modularity is most distinct toward the bases of food chains, and the four modules identified in the community are based primarily on the differential utilization of basal and low trophic level resources. High trophic level predators, however, such as the Caribbean reef shark, feed across modules, both uniting modules via top-down effects, and reducing the modularity of the system. Furthermore, many of those predatory species are involved in very strong biotic interactions [79]. The apparent modularity of the modern community might therefore be an anthropogenically amplified effect, because many high trophic level predators have been either extirpated from reef communities throughout the Caribbean, or are now present in very low numbers. The fossilized food web retains the modularity of the modern web, perhaps because it too lacks many higher trophic level species, though for preservational reasons. This emphasizes the caution which should be exercised in the analysis of modern, post-disturbance communities, and it is useful to speculate that assessments of reef community modularity based on historical and sub-fossil records [37] could yield significantly different results. It is also worth considering, in this context, whether the introduction of the invasive lionfish, *Pterois volitans,* into the Caribbean has restored some of the effects of predation now lacking in those communities.

We tested our ability to recreate a coral reef food web from fossil data only, by assuming the simulated fossil reef as a starting point. Following procedures developed for terrestrial paleocommunities [67], trophic guild structure was used as the basis to aggregate species into biotically interacting units. We used a heuristic approach to first identify trophic guilds in the modern food web. If we limit our ability to recognize guilds according to the limits of fossil preservation, we find a total of 42 guilds, whereas the maximum number identified in the modern community is 149. Nevertheless, Simpson (1-D) and Shannon (H) Indexes for modern and fossil guild diversity were almost identical despite a loss of guilds in the fossil community.

Finally, species level food webs were reconstructed from the fossil and guild data using stochastic techniques described in [59]. One reconstruction, based on the indegree distribution of the modern web, fails to recreate the trophic level structure, and hence hierarchical arrangement of the modern community. In contrast, a second reconstruction, based on a model in-degree distribution developed in previous studies [67], creates a trophic level and hierarchical structure statistically indistinguishable from the modern community. The greater success of the latter in-degree distribution is a consequence of its overestimation of species dietary breadths, thereby compensating for losses incurred during fossilization. The important implication of this successful recreation is that coral reef communities can indeed be recreated from fossil data, and subjected to the types of dynamic analyses essential for forecasting the behavior of those communities under broad magnitudes of environmental and biotic disturbance. This emphasizes the validity and usefulness of paleoecological data to the field of conservation biology. The creation of deep-time paleocommunity food webs has the ability to enrich and advance our current knowledge of how natural systems behave, especially in response to future environmental changes.

## Acknowledgements

We wish to thank Courtney Chin, Chrissy Garcia and Rhiannon Roopnarine for the food chain illustrations and other graphical assistance. We also thank the CAS Paleoecology Discussion Group, including Allen Weik, for many useful discussions which contributed to this chapter. Finally, thanks to Kenneth Angielczyk and Carol Tang for asking difficult questions.

## Appendix 1

### Hypergeometrie variance

The hypergeometrie distribution describes the probability that, given a population with a featured subset of observations, and a sample from the population, that a random selection of individuals from the population will include a certain fraction of the sample comprising individuals with the featured characteristic. In this case, the population consists of 728 species (*N*), of which 433 are featured as fossilized (*K*). A sample is the richness of a trophospecies (*n*), of which a subset are observed as fossilized (*k*). The expected fossilization, or hypergeometric mean value is

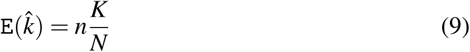

The variance of the expectation is given as

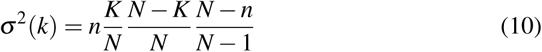

This value grows initially as *n,* because the number of unique ways in which k objects may be selected also grows. The value declines, however, as *n* → *N* (Fig. 12).

## Appendix 2

**Fig. 12.**
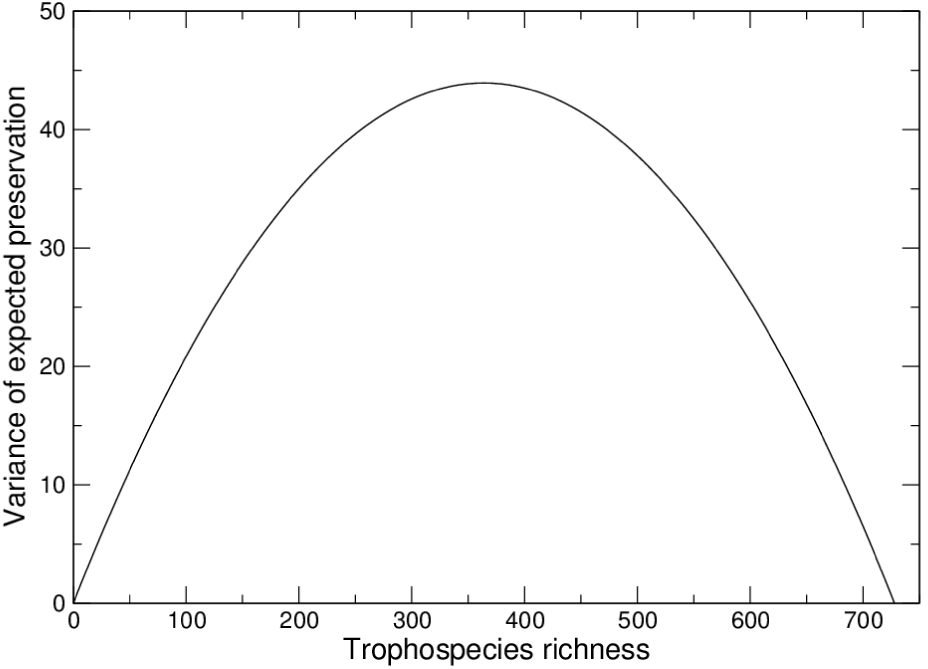
Hypergeometric variance given a total population of 728 species, and trophospecies of increasing size.

**Table 1.** List of guilds recovered at the 0.6 threshold of interaction overlap. Modern and fossil guild richnesses are shown. Guilds containing single taxa have those taxa listed by taxonomic or unique common name. Fossil guild richnesses with an asterix * were estimated.

| Guilds | Modern | Fossil |
| --- | --- | --- |
| Bacteria | – | – |
| Phytoplankton | 29 | 15 |
| Nanno-zooplankton | 18 | 3 |
| Macrophytes, diatoms | 28 | 14 |
| Sponges | 53 | 31* |
| Corals | 70 | 64 |
| Micro-detritivores | 19 | 18 |
| Corallivorous polychaetes | 2 | 0 |
| Zooplanktonv | 44 | 45* |
| Gorgonians | 15 | 0 |
| Benthic carnivores | 22 | 0 |
| Eucidarid echinoids | 1 | 1 |
| Hard benthic macroinvertebrates | 123 | 90 |
| Cypraeids | 3 | 3 |
| *Sinum* | 1 | 1 |
| *Charonia* | 1 | 1 |
| Epiphyte-grazing gastropods | 3 | 3 |
| Bryozoans | 25 | 20 |
| Endolithic polychaetes | 3 | 3 |
| Soft benthic macroinvertebrates | 80 | 75 |
| Polychaete consuming gastropods | 6 | 6 |
| Molluscivorous crustaceans | 1 | 1 |
| Macroinvertebrate predators I | 60 | 25 |
| Stomatopods I | 2 | 2 |
| Large grazers of macroalgae and invertebrates | 79 | 47 |
| Macroinvertebrate predators II | 4 | 4 |
| *Octopus* | 1 | 0 |
| Macroinvertebrate predators III | 11 | 2 |
| Stomatopods II | 3 | 3 |
| *Hemiramphus brasiliensis* | 1 | 0 |
| *Trachinotus goodei* | 1 | 0 |
| Pelagic piscivores I | 8 | 8 |
| *Mycteroperca bonaci* | 1 | 1 |
| *Gymnothorax moringa* | 1 | 0 |
| *Lutjanus analis* | 1 | 0 |
| *Scomberomorus cavalla* | 1 | 1 |
| Green sea turtle | 1 | 1 |
| Hawksbill turtle | 1 | 1 |
| Loggerhead turtle | 1 | 1 |
| Blacktip shark | 1 | 1 |
| Oceanic whitetip shark | 1 | 1 |
| Caribbean reef shark | 1 | 1 |

